# BASE: a novel workflow to integrate non-ubiquitous genes in comparative genomics analyses for selection

**DOI:** 10.1101/2020.11.04.367789

**Authors:** Giobbe Forni, Angelo Alberto Ruggeri, Giovanni Piccinini, Andrea Luchetti

## Abstract

Inferring the selective forces that different ortholog genes underwent across different lineages can make us understand the evolutionary processes which shaped their extant diversity. The more widespread metric to estimate coding sequences selection regimes across across their sites and species phylogeny is the ratio of nonsynonymous to synonymous substitutions (dN/dS, also known as *ω*). Nowadays, modern sequencing technologies and the large amount of already available sequence data allow the retrieval of thousands of genes orthology groups across large numbers of species. Nonetheless, the tools available to explore selection regimes are not designed to automatically process all orthogroups and practical usage is often restricted to those consisting of single-copy genes which are ubiquitous across the species considered (*i.e.* the subset of genes which is shared by all the species considered). This approach limits the scale of the analysis to a fraction of single-copy genes, which can be as lower as an order of magnitude in respect to non-ubiquitous ones (*i.e.* those which are not present across all the species considered). Here we present a workflow named BASE that - leveraging the CodeML framework - ease the inference and interpretation of selection regimes in the context of comparative genomics. Although a number of bioinformatics tools have already been developed to facilitate this kind of analyses, BASE is the first to be specifically designed to ease the integration of non-ubiquitous genes orthogroups. The workflow - along with all the relevant documentation - is available at github.com/for-giobbe/BASE.

## Introduction

Selection can drive the evolution of genes by constraining changes in their sequences (purifying selection) or by favoring new adaptive variants (positive selection). Quantifying its mode and strength is a key step to understand the diverse evolutionary histories of ortholog genes across different clades. Statistical models of molecular evolution have proved as a fundamental approach to investigate such processes and can be divided into those based on comparing divergence and segregating polymorphism - such as the MK test and its extensions (McDonald and Kreitman 1991) - and those based on multi-species sequence divergence - also known as codon models. The two approaches use different conceptual frameworks and are better applied for analyses at different timescales, with the first approaches more suited to investigate recent processes and the latter ones more apt to infer older events (Mugal et al 2014).

Approaches based on the sequence divergence among multiple species are cornerstones in the estimations of patterns of sequence evolution and selection regimes. After the first models were developed to infer the strength of selection on protein-coding sequences globally across their sites and species phylogeny (Goldman and Yang, 1994; Muse and Gaut, 1994), subsequent elaborations allowed for variation across lineages (Yang, 1998), sites (Nielsen and Yang, 1998; Yang et al. 2000; Anisimova et al. 2001) and both (Yang and Nielsen 2002; Zhang et al. 2005). Pairwise comparisons between models can be performed using likelihood-ratio tests to understand which one better reflects the molecular evolution of a group of ortholog genes (Anisimova et al. 2001). The interpretation of all these models is largely based on the dN/dS parameter (Kimura, 1977; also known as ω), which consists in the ratio of nonsynonymous substitution rates (non-synonymous mutations over non-synonymous sites; dN) to synonymous substitution rates (synonymous mutations over synonymous sites; dS). This metric is fundamental to investigate the extent to which selection modulates sequence evolution of the protein-coding portions of genes. While d*S* are assumed to evolve neutrally, d*N* are expected to be exposed to selection, as they change the aminoacidic structure of proteins. Despite some of these assumption have been challenged (Davydov et al., 2019; He et al., 2020), analyses based on codon models have proved themselves as key approaches in comparative genomics, such as investigating positive selection connected to evolutionary innovations (Parker et al. 2013; Zhang et al. 2014; Li et al. 2014) or testing the relaxation of selective constraints after trait decay (Liu et al. 2019; Policarpo et al. 2020). In other instances, these approaches have been used to observe genome-wide effects linked to events such as shifts in environmental niches or the loss of recombination in asexual genomes (Plazzi et al., 2017; Bast et al. 2018).

Several pieces of software have been developed to infer codon models for coding sequences: Selecton (Stern et al. 2007), HyPhy (Pond et al., 2005), TreeSAAP (Woolley et al 2003) and the CodeML program in the PAML package (Yang, 2007). The latter program was also subject to several implementations, such as IDEA (Egan et al., 2008), PAMLX (Xu and Yang 2013), SlimCodeML (Valle et al 2014), IMPACT_S (Maldonado et al. 2014), LMAP (Maldonado et al. 2016), ete-evol in the ETE3 package (Huerta-Cepas et al. 2016), VESPA (Webb et al. 2017), BlastPhyMe (Schott et al. 2019) and EasyCodeML (Gao et al., 2019). With the increment of genomics and transcriptomics studies, it has become rather common to analyze thousands of genes for up to hundreds of species and - despite CodeML still appearing to be the most widely used piece of software - all of its aforementioned implementations try to increment its ease of use in the context of comparative genomics.

Our focus developing BASE has been mainly directed to facilitate the integration of an often overlooked - yet incredibly large - portion of genomes. Orthogroups (OGs) can differ in many aspects, such as the inclusion of single-copy or multi-copy genes. OGs can also be made up of ubiquitous (*i.e.* the subset of genes which is shared by all species considered) or non-ubiquitous (*i.e.* those which are lacking in some of the species considered) genes. In comparative genomics datasets, the majority of single-copy genes OGs consists of non-ubiquitous genes: as an exploratory example, we analyzed 20 recent datasets and found that - among single-copy genes OGs - the average proportion of those including non-ubiquitous genes is 73.4% (**Fig. 1**). Non-ubiquitous genes OGs are mostly overlooked in selection analyses - which are typically based only on single copy and ubiquitous genes - due to the lack of automated approaches for their inclusion. Nonetheless, disregarding such a large portion of genes may potentially conceal important evolutionary processes and therefore we developed a novel workflow specifically for this purpose.

**Fig. 1.**
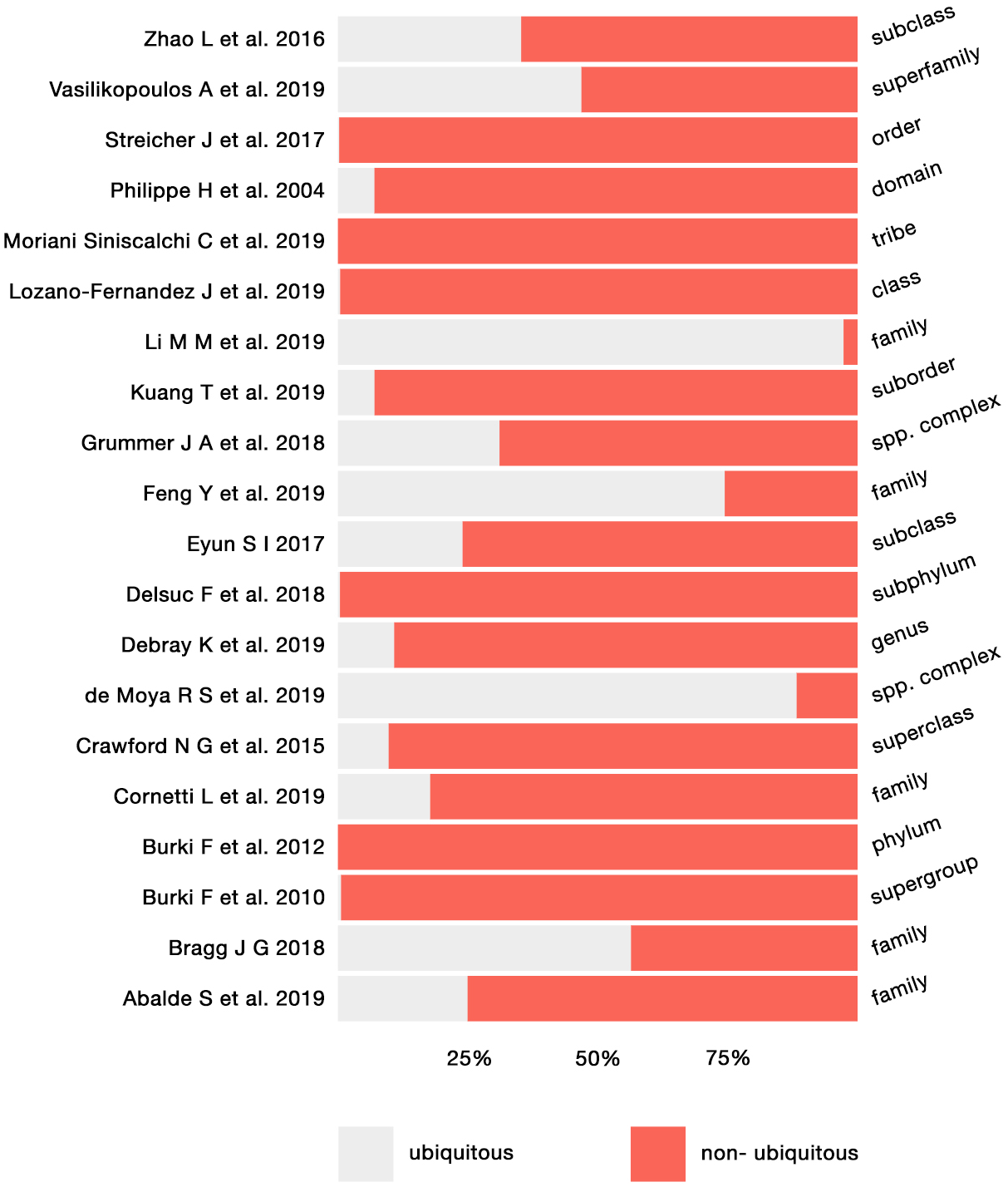
The proportion of ubiquitous and non-ubiquitous genes was calculated in 20 datasets, varying in the taxonomic level considered (from family to phylum). The average percentage of non-ubiquitous genes is 73,36 % while for ubiquitous ones is 26,6%.

## Implementation

The BASE workflow is written in BASH and R and has been tested on Linux operating systems, such as centOS 8. As it extensively leverages GNU utilities, its usage is restricted to Linux distributions. It consists of two main steps: in the first one (“analyze”), evolutionary model parameters are inferred across alignment sites and tree branches for the different OGs, while the subsequent step (“extract”) allows to retrieve the different metrics associated to specific branches or clades in the species tree. CodeML provides the statistical and computational framework to perform these analyses and is at the core of the workflow, whose general description is reported in **Fig. 2**.

**Fig. 2.**
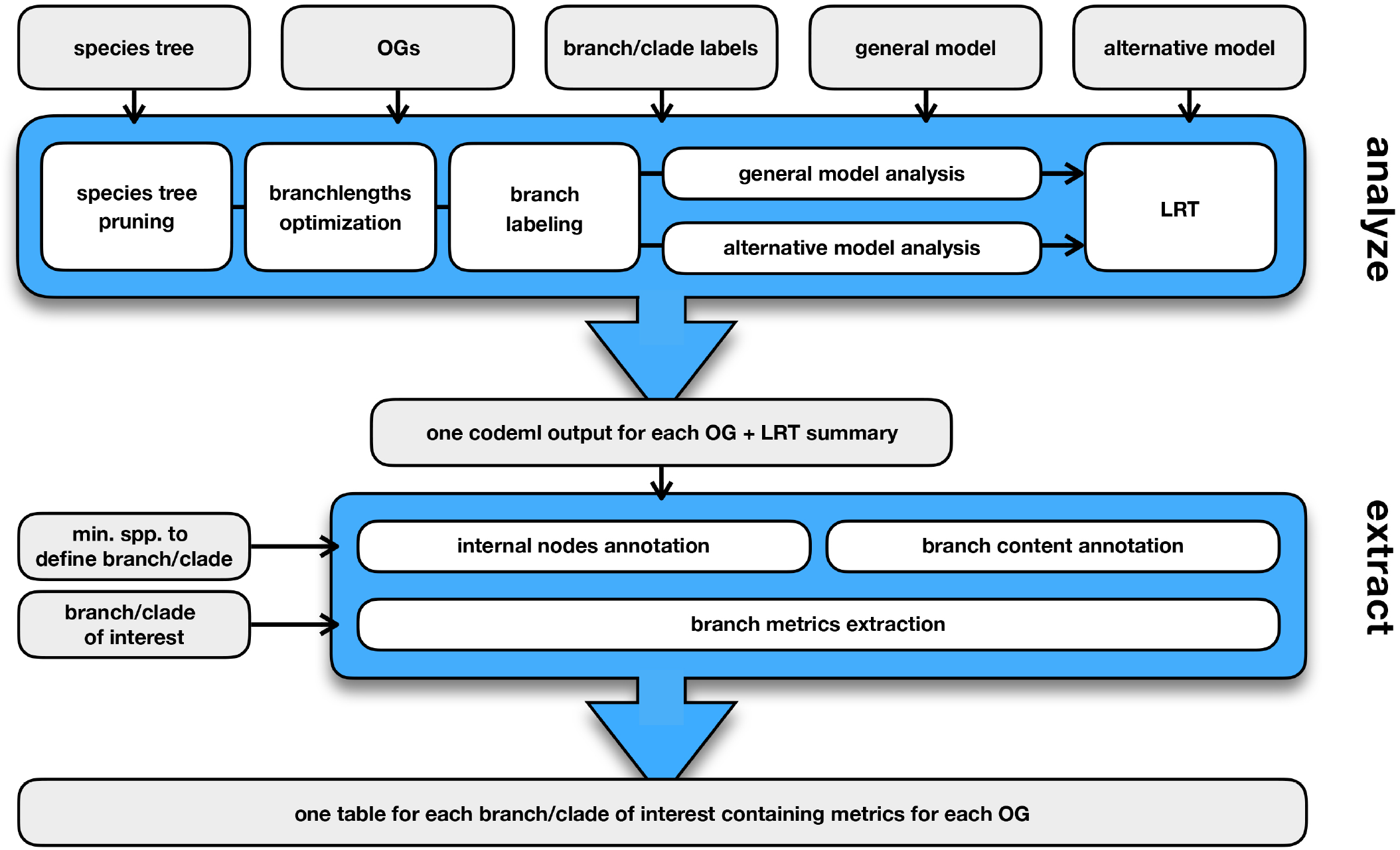
BASE workflow consists of the two steps: “analyze” and “extract”. The two steps can be carried out independently and CodeML output generated outside of the workflow can be processed in the “extract” step.

The inputs required to start an analysis are: a) a species tree in the Newick format; the tree can be multifurcating but has to include all the species present across the different OGs; b) a folder containing aligned OGs, in fasta format with headers matching exactly with the species names found in the species tree; c) two CodeML control files describing two nested models - where one is a specific case of the other; all the parameters in control files can be customized, with branch-site, branch and site models all supported; d) a file including the branches or clades in the species tree for which the different metrics will be retrieved, defined by their associated species.

In the “analyze” step, the workflow initally checks the alignments for the presence of stop codons using transeq of the EMBOSS package (Rice et al. 2000), on the basis of the genetic code specified in the control files; genes which include stop codons are excluded from subsequent analyses. Branch length optimization is then carried out for each OG using RAxML (Stamatakis 2014) with a codon-aware GTR substitutions model; subsequently the two CodeML runs configured with the general and the alternative models are performed and compared through a Likelihood Ratio Test (LRT) using R (R Core Team 2013). LRTs results are summarized in a table and the output relative to the best-fit model is selected for each OG. The default behavior of BASE is to process all OGs, whether consisting of ubiquitous or non-ubiquitous genes, but the user can limit the analysis to just ubiquitous ones. When the analysis is configured to consider also OGs of non-ubiquitous genes, the species tree will be pruned on the basis of the species present in each OG using ape R package (Paradis et al. 2004), prior to branch length optimization. It is possible to label specific branches or clades of the phylogenetic tree: to do so a file listing all the species associated to a clade need to be specified, so that the relative tags will be added to the tree using phangorn R package (Schliep 2011).

The “extract” step can be carried out subsequently to the “analyze” one, using as input the folder generated by the previous step; otherwise CodeML outputs generated by means other than BASE can be used as well. In a first place this step will annotate internal nodes of each OG tree to match the output of CodeML and will list all species associated to each branch of the phylogeny. Subsequently, the pipeline will create a table for each branch/clade specified by the user, containing the dN/dS, dN, and dS values relative to the best-fit model for each OG.

Equivalent branches/clades can be identified in a phylogeny even in the absence of some species (**Fig. 3**). For example, a clade - and its stem branch - made up by tens of species can be considered to be still present if we subtract a few species from the phylogeny, either in the ingroup or in the outgroup. In BASE “extract” step it is possible to include non-ubiquitous genes OGs with two approaches: (a) including OGs of non-ubiquitous genes just relatively to an outgroup - *i.e.* species external to the clade(s) of interest; (2) include OGs of non-ubiquitous genes also relatively to an ingroup - *i.e.* clade(s) of interest. In the latter scenario, the user can configure a cutoff of missing species relative to the branches/clades of interest, which can be specified by either an absolute number or a proportion (for example: if 0.8 is specified, at least the 80% of the clade’s species need to be present in a given OG in order to include it in the analysis). If representatives of a given clade (or its stem branch) do not meet the selected criteria, this will be stated in the final output and no associated metrics will be reported.

**Fig. 3.**
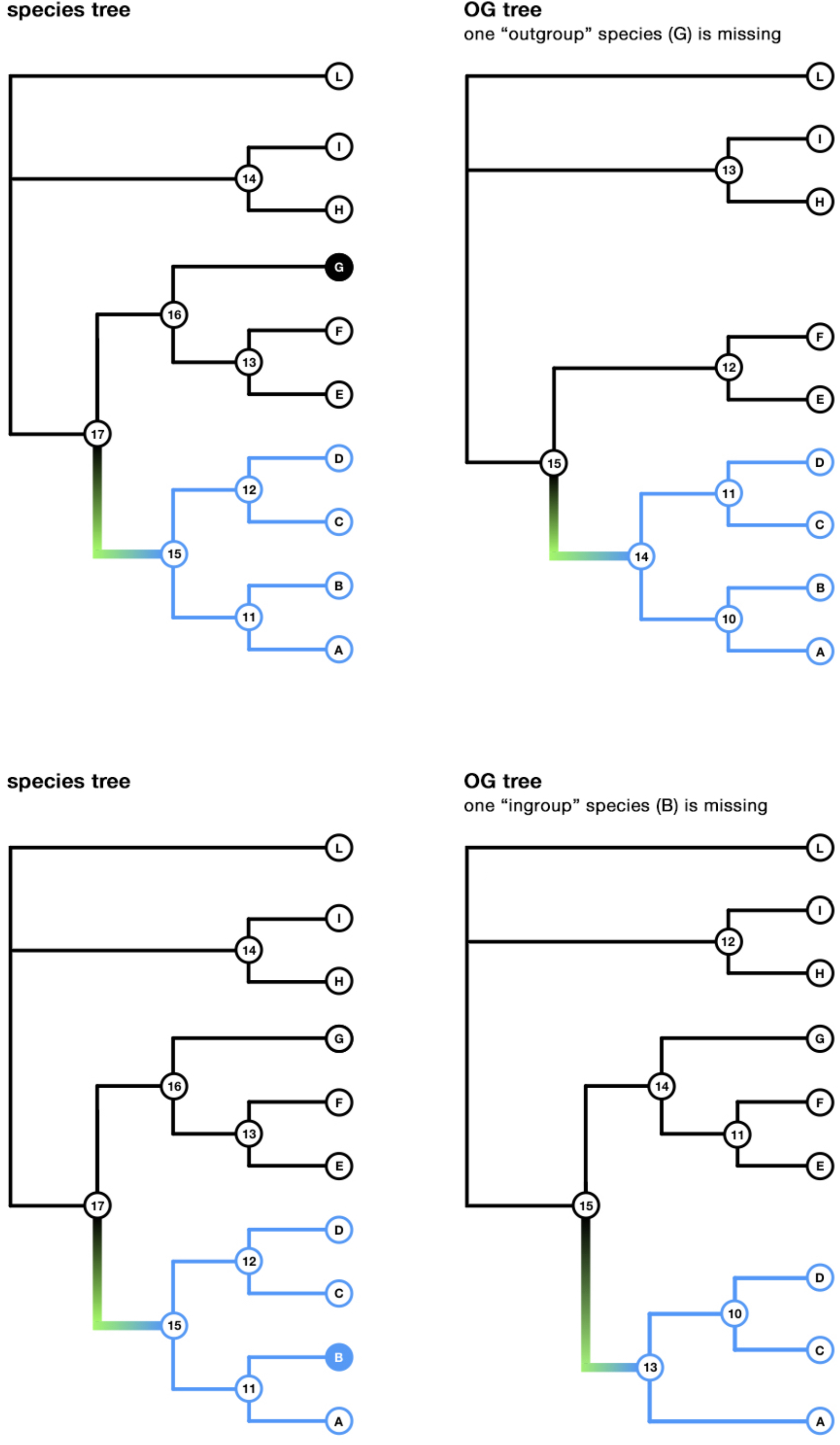
Trees of non-ubiquitous genes OGs differ in their structure from the species tree. Other than restricting the analysis to ubiquitous genes, BASE can: (a) include OGs of non-ubiquitous genes just relatively to a clade of interest (outgroup; top figure); (b) include OGs of non-ubiquitous genes also relatively to a clade of interest, on the basis of a user-defined threshold (ingroup; bottom figure). In the example the branch leading to a clade of interest is highlighted in green, but the same applies when whole clades are considered.

Even if the focal feature is the integration of single-copy non-ubiquitous genes OGs into selection analyses, BASE provides other features that ease the inference, comparison and interpretation of codon models. Labeling and/or retrieving model parameters for specific branches/clades in large phylogenies can be a tiresome process, which is instead made easy by the approach implemented in our workflow. BASE can also process batches of OGs simultaneously - substantially cutting down processing times - and implements a large number of error-messages which can definitively ease the user experience.

## Conclusions

BASE is a workflow for analyses on selection regimes that integrates several popular pieces of software, with CodeML at its core. It has been conceived to ease the integration of non-ubiquitous genes OGs into comparative genomics analyses for selection, yet it implements many other features and quality-of-life improvements. BASE can represent a useful tool to address theoretical and biological questions, such as the impact of sampling on dN/dS estimates or whether ubiquitous genes tend to have different evolutionary constraints with respect to non-ubiquitous ones. We hope that our effort proves to be a useful tool for studying molecular evolution and that it generates some interest towards the integration of non-ubiquitous genes OGs in selection analyses. BASE is an ongoing project, and we welcome bug reports, feedback and suggestions for feature implementations.

All the documentation, including detailed tutorials to explore BASE functionality can be found at github.com/for-giobbe/BASE.

## Author contributions

G.F. conceived the idea; All authors implemented ideas and design; G.F. led the writing of the manuscript and on-line resources; All authors contributed to the drafts and gave final approval for publication.

